# An automated barcode tracking system for behavioural studies in birds

**DOI:** 10.1101/201590

**Authors:** Gustavo’, Jacob M. Graving, James A. Klarevas-Irby, Adriana A. Maldonado-Chaparro, Inger Mueller, Damien R. Farine

## Abstract

1. Recent advances in technology allow researchers to automate the measurement of animal behaviour. These methods have multiple advantages over direct observations and manual data input as they reduce bias related to human perception and fatigue, and deliver more extensive and complete data sets that enhance statistical power. One major challenge that automation can overcome is the observation of many individuals at once, enabling whole-group or whole-population tracking.
2. We provide a detailed description for implementing an automated system for tracking birds. Our system uses printed, machine-readable codes mounted on backpacks. This simple, yet robust, tagging system can be used simultaneously on multiple individuals to provide data on bird identity, position and directionality. Further, because our codes and backpacks are printed on paper, they are very lightweight.
3. We describe the implementation of this automated system on two flocks of zebra finches. We test different camera options, and describe their advantages and disadvantages. We show that our method is reliable, relatively easy to implement and monitor, and with proper handling, has proved to be safe for the birds over long periods of time. Further, we highlight how using single-board computers to control the frequency and duration of image capture makes this system affordable, flexible, and adaptable to a range of study systems.
4. The ability to automate the measurement of individual positions has the potential to significantly increase the power of both observational and experimental studies. The system can capture both detailed interactions (using video recordings) and repeated observations (e.g. once per second for the entire day) of individuals over long timescales (months or potentially years). This approach opens the door to tracking life-long relationships among individuals, while also capturing fine-scale differences in behaviour.

## Introduction

Studying behaviour is central to addressing a broad range of research questions in the fields of neurobiology, ecology, and evolutionary biology. Nevertheless, collecting accurate and complete behavioural data remains a challenging task (Crall *et al.* 2015). Although direct observation is still an important method for gathering data, a variety of automated methods are now frequently used to accelerate data collection and reduce the effects of human intervention. Video recording has become common practice for studying both captive (Togasaki *et al.* 2005; Nagy *et al.* 2013; Perez-Escudero *et al.* 2014; Rojas Mora *et al.* 2014; Ihle *et al.* 2015) and wild organisms (Togasaki *et al.* 2005; Scheibe *et al.* 2008). However, manually measuring behaviour from images, like photos and videos, is extremely time consuming and may still have the same limitations as direct observation, including human bias and fatigue. Manually identifying individuals is also challenging, which limits the use of this approach to species with individually-distinct features (Perez-Escudero *et al.* 2014). Recent advances in automated, image-based tracking methods have solved these issues in a variety of ways. Unfortunately, many of these solutions rely on complex, computationally-intense algorithms, often require keeping animals in simplistic, unnatural environments, and may not reliably preserve identities over long periods of time or across contexts (e.g. Perez-Escudero *et al.* 2014). One alternative, which has been explored in a few recent studies (Mersch *et al.* 2013; Nagy *et al.* 2013) is to fit machine-readable tags to individuals, allowing for faster, more reliable tracking. This method offers exciting new opportunities, such as studying social behaviour in complex, naturalistic environments over long timescales, and across multiple experimental conditions. Here we provide details of how to implement such a system for songbirds.

The development of methods for tracking individuals plays an important role in our ability to study animals. In addition to the limitations of human observers to process multiple streams of information simultaneously (such as the actions of several individuals in a group), many studies still rely on using relatively small datasets to estimate broad patterns. One example is the use of focal follows, where a single individual is tracked for a period and all of its interactions with others are recorded. While doing so, all the interactions among others are not recorded. This means that even with very intensive monitoring, the maximum number of dyadic observations that can be made are N-1, where N is the number of individuals present. Sparseness in the resulting datasets can impact the ability to successfully test hypotheses (Farine & Strandburg-Peshkin 2015). Further, these studies can also suffer from temporal autocorrelation (most data on a focal is collected within a short period of time; Whitehead 2008). Studies that cannot extract data with sufficient resolution also lead to concerns about the use of animals in research if they cannot robustly test the hypothesis, as poor data collection can heighten the rates of true and false positives.

Several technologies enable more detailed tracking of individuals than what is possible by human followers. An increasingly common method for tracking small birds is Passive Integrated Transponder (PIT) tags (Boarman *et al.* 1998). These small microchips generate a disturbance in the electric field of Radio Frequency Identification (RFID) antennas, and the pattern of disturbance can be used to encode a unique identity for each tag. Because PIT tags require no battery power, they enable large-scale deployment over long periods of time and have been used in both laboratory (Griffith *et al.* 2010; Weissbrod *et al.* 2013; Boogert *et al.* 2014; Farine *et al.* 2015) and field conditions (Broderick & Godley 1999; Bonter & Bridge 2011; Mariette *et al.* 2011; Steinmeyer *et al.* 2013; Farine *et al.* 2014; Adelman *et al.* 2015; Aplin *et al.* 2015; König *et al.* 2015). However, two major limitations of this technology are that (1) if multiple tags are in the antenna field, none are detected (hence no simultaneous detections are possible); and (2) detections are confined to very small areas where antennas are present. As video hardware and computer vision software have improved, one popular alternative is to implement automated, image-based tracking of animals (Dell *et al.* 2014; Perez-Escudero *et al.* 2014; Jolles *et al.* 2015; Rosenthal *et al.* 2015). Great efforts have been made to overcome the challenge of tracking multiple animals while maintaining individual identities. Some algorithms can identify individuals using their distinct patterns (Berger-Wolf *et al.* 2015) and even maintain individual identities by recognizing subtle differences in colouration (Perez-Escudero *et al.* 2014). Unfortunately, these methods require animals with distinguishable features, or keeping animals in highly artificial conditions with a simple background and uniform lighting. Moreover, tracking approaches typically require high-frequency video, which collects much more data than what is often required, and introduces significant hardware costs in terms of video quality, processing and storage.

One solution for identifying individuals is to attach a machine-recognisable marker to each animal. Studies on social insects were the first to implement 2D barcodes (hereafter barcodes) (Mersch *et al.* 2013; Crall *et al.* 2015) with a unique pattern of black and white squares that can be identified and matched to a library of known codes. Insects have been good models for using such technology because these barcodes can be directly glued onto their bodies, and they can be applied to hundreds of individuals simultaneously because codes are inexpensive to make (using only waterproof paper). Similar approaches have been used on fish and birds, but these typically involved simpler tracking of a coloured tag temporarily fitted to individuals (eg. Nagy *et al.* 2013), and few details are available on their implementation. Despite representing a major advance in data quality, barcodes are rarely used over long periods of time and in semi-natural conditions. This is especially surprising when considering that tags can be implemented for very little cost and tailored to suit a range of experimental conditions.

Both in captivity and in the wild, birds represent a challenge for automated tracking because they often lack markings that allow for identification of individuals (because feathers move). Further, many birds are highly social, but current data collection methods, specifically PIT tags, can only detect single individuals present at focal locations, such as nest boxes (Schlicht *et al.* 2015; Santema *et al.* 2017), feeders (Firth *et al.* 2016), or puzzle-boxes (Aplin *et al.* 2015). To cope with issues of observing multiple individuals, as well as to overcome the limitations of human observers, we developed a barcode tracking system for birds. Such a system identifies—and allows for tracking of—individuals’ positions and orientations over time. Here we describe the design and deployment procedures for backpack-mounted barcodes, as well as the required monitoring and maintenance of the system over long periods of time, to assure bird safety and reliable data collection. We discuss the materials used, different camera systems for capturing image data, and other considerations associated with data collection. We provide details on the process of extracting data from the images, and what software is available for this purpose. Finally, we highlight potential behaviours that can be measured using such a system and possible applications in further studies.

## MATERIALS AND METHODS

### Study population

We tested our barcode tracking system on domesticated zebra finches (*Taeniopygia guttata*). The zebra finch is a model species widely used in behavioural studies (David *et al.* 2011; Schuett *et al.* 2011; Mariette & Griffith 2012; Boogert *et al.* 2014; Ruploh *et al.* 2014; Farine *et al.* 2015; Wuerz & Kruger 2015; Kriengwatana *et al.* 2016). They are social birds, living in colonies of 50–100 individuals (Zann 1994), and in captivity can be kept in large groups. This makes them a suitable organism to test our tracking system.

We tested our system on two flocks of domesticated zebra finches, held in separate indoor aviaries in the Max Planck Institute for Ornithology in Radolfzell, Germany, with indoor aviary lighting matching the local day/night patterns. Each flock was held in a 2 × 2 × 2-m metal-mesh cage and provided with a complex arrangement of natural branches, feeders, drinking water, a bathing tray and wood chips as floor cover. We supplied both millet seeds and water ad libitum, except during food-based assays (see below). No nesting material or nest boxes were available during the length of our trials to prevent the birds from breeding. Each flock consisted of 28 adult individuals in 1:1 sex ratio. We tested prototype backpacks in each flock from September to November 2016. From December 2016 through to the end of March 2017, we fit backpacks to all members of the flocks (except those that could not take a backpack, see below). Birds were therefore fitted with individual backpacks for up to 4 months, with some individuals carrying backpacks continuously over a period up to 7 months. Each bird was also fitted with leg-bands for identification, consisting of a numbered closed metal band and two plastic bands in a colour combination that was unique in each aviary. This study was conducted under Ethics Permit 35-9185.81/G16/73 issued by the state of Baden-Württemberg.

### Barcode tracking system

The barcode tracking system consists of 3 components: (1) a backpack fitted with a barcode; (2) recording device(s), and (3) processing software and hardware. In this section, we describe the design of the backpack (i.e. structure carrying the barcode), its fitting procedure (i.e. deployment), and the monitoring and maintenance of the codes.

#### Backpack design

Backpacks consisted of three main parts: the backpack structure and tag mount, the tray, and the straps (Fig. 1). We constructed the structure using 70 × 10-mm strips of waterproof and tearproof paper (Xerox^®^ -Premium Never Tear- 95μm). We built this structure by laser printing templates on an A4 sheet of paper (Fig. 1A, template provided in Supplemental Materials). Each template was cut out, folded, and glued into a loop to form the tag mount (Fig. 1B, 1F), which provided a raised surface to keep the barcode above the feathers. We created a 3D-printed black plastic tray (Fig. 1C) that housed a barcode that was printed on the same type of paper as the backpack structure (Fig. 1D). This barcode was glued into the recess of the plastic tray. The black plastic is an important feature as it reinforces the border that frames the barcode and prevents the birds from damaging the edges, which makes the code unreadable by the software. This tray also keeps the code flat, rigid and visible to the cameras. We glued this tray with the code onto the backpack mount (Fig. 1F). Although a well deployed backpack should keep this tray behind the wing joints, we rounded the external corners of the tray (Fig. 1C) to prevent injuries and wing rubbing.

We narrowed the front strip of paper to fit between the scapulae of the bird, and punched four round holes (^~^ 1mm diameter, Fig. 1E) to attach the elastic string that formed the straps of the backpack around the bird (Fig. 2). For each backpack, we used a single piece of string 25 cm long which we looped through the rear holes on the paper, crossed under the backpack, tied on the front holes, and kept the leads loose to allow for individual adjustment during deployment. For zebra finches, we used a 28 × 6mm front strip and a 10 × 10-mm mount raised 6 mm.

**Figure 1.**
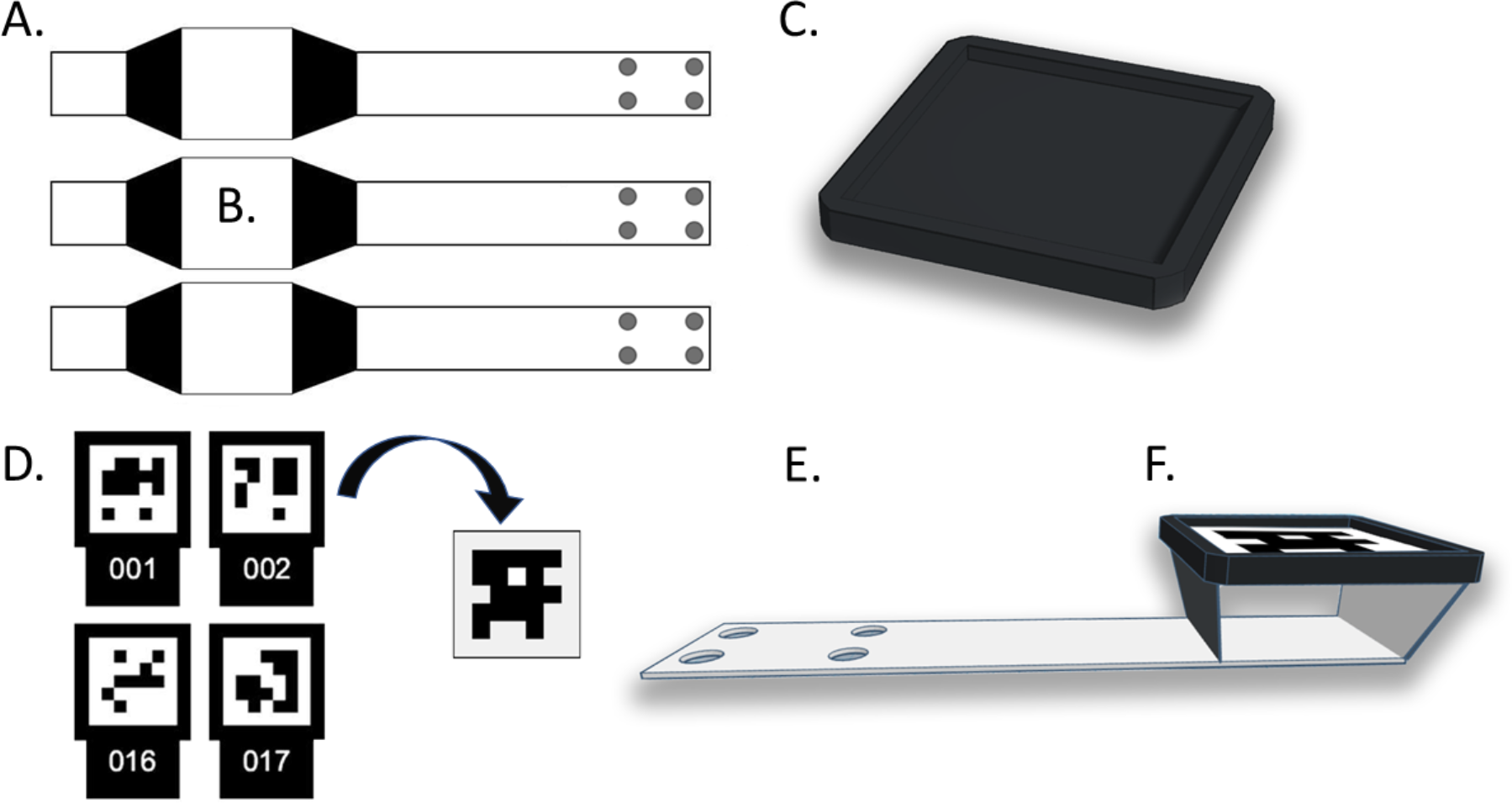
Components and assembly of the backpack-mounted barcodes. (**A)** Template layout. (**B)** Mount area where plastic trays (**C**) are glued. (**D)** Barcode layout and cutout to be glued on the tray. (**E)** Assembled backpack with tray and code raised on the mount (**F**).

#### Backpack deployment

The general procedure for fitting backpacks was as follows: (1) we caught and measured each bird, and recorded its health status; (2) we fit the backpack; (3) briefly placed each bird in a small observation cage; (4) released and monitored each bird in their permanent housing, and (5) performed periodic health checks.

Once we confirmed the birds were in good health (step 1), we fit a completely assembled backpack to each bird (step 2). We pre-tied the string on the backpack with a simple slipknot and then pulled the straps over the bird’s head until the front strip sat on the interscapular area, carefully pulling each wing through the string straps. We found that the best fit was achieved when the leading edge of the raised section was below the elbow joint of the wing, and the trailing edge was above the rump (Fig. 2). Once the backpack was in its final position,we tightened the string around the body, adjusting according to the size of each bird. The tightness must be firm enough to hold the backpack in position while preventing the bird to put its feet/toes inside of the loop, but also loose enough to allow the birds to fly and move freely and to avoid blocking the crop. The front strip and the string loops were covered by the feathers, while only the mount with the plastic tray and the barcode are visible. The mount must be positioned behind all the wing bones and joints, where only feathers can be in contact with it.

After fitting the backpack to a bird, we briefly placed the bird in a small observation cage (step 3). This step was critical for evaluating each individual’s behaviour to assure the backpack was not interfering with normal movement. Most birds tried to pull the backpacks or the straps off during this period. In our experience, the intensity and duration of this behaviour was not necessarily a signal of an ill-fitting backpack and, on the contrary, it helped to accommodate all the new elements. A well-fitted backpack allows the bird to move freely, with minimum or no interference for flying, walking, landing or perching. Once birds performed these without difficulty, we made a final check of the adjustments, tightened the knot near the neck of the bird, secured the knots with cyanoacrylate glue, and cut any excess string from the leads. We also trimmed some coverts around the mount to prevent any obstructions on the codes that might hinder readability. We found that this acclimation process worked better when birds were kept in small groups (2-5) and in a different room to us, as it reduced stress and allowed for allopreening, feeding and undisturbed movement.

**Figure 2.**
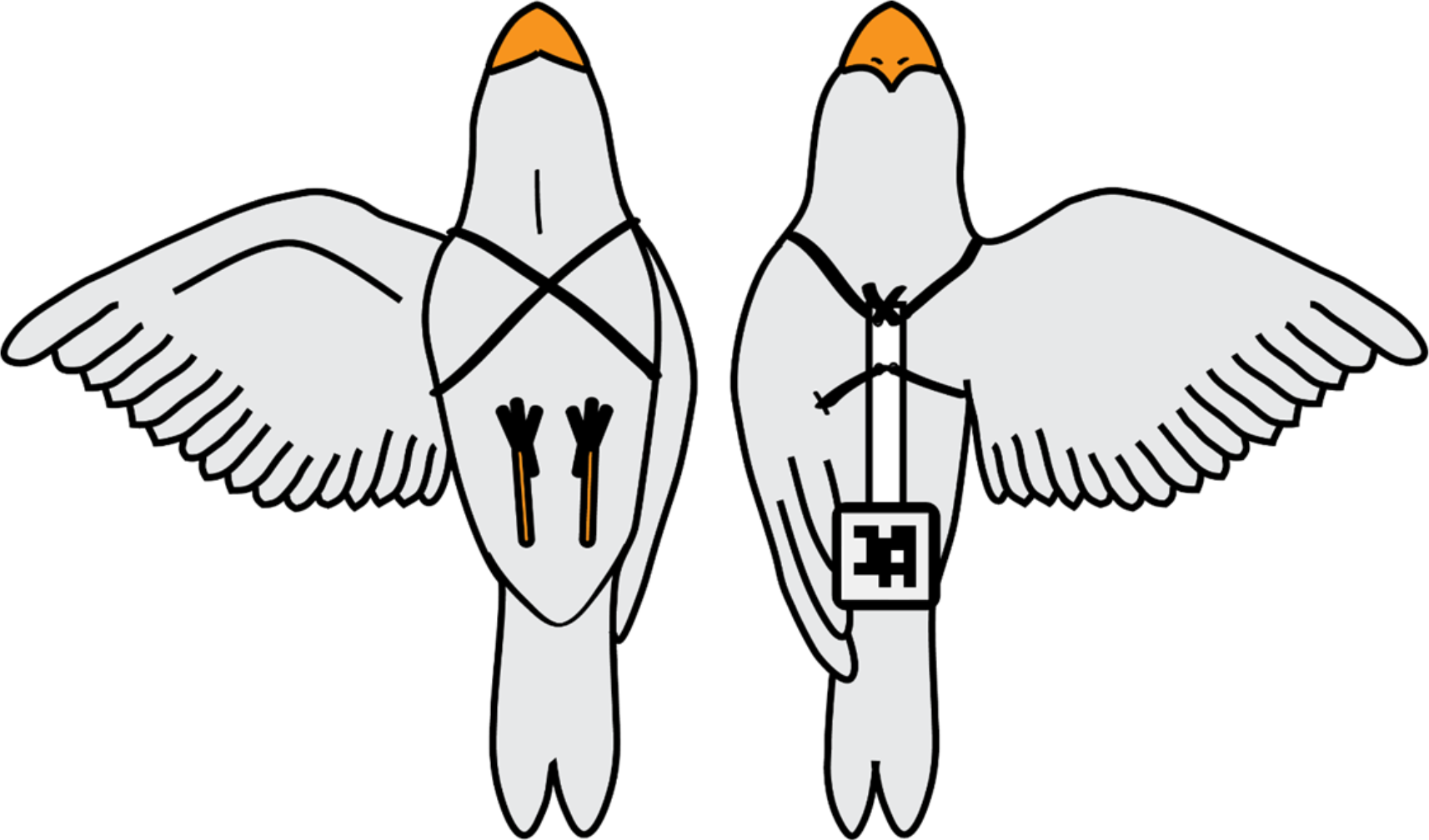
Left: bottom view of bird with backpack. The string sits in front and behind the wings and crosses on the chest of the bird. Right: Top view. The string is tied at the anterior end of the backpack which goes between the scapulae, and the mount with the barcode sits on the rump, behind the wing joints.

Every time we observed a bird with hindered movement or unusual behavior possibly related to the backpack, we checked and re-adjusted the straps. In some cases, we completely removed the backpack, let the birds rest to reduce stress, and observed their behaviour without the backpack before trying another deployment. A few birds (4 of 58) could not be tagged properly despite having a well-fitted backpack and appearing to be in good health. We removed these subjects from the fitting procedure.

#### Backpack Monitoring

We monitored the birds regularly, either during our experiments or during care-taking activities, and constantly looked for unusual behaviour. This monitoring is important to prevent injuries or to detect early symptoms of health issues, either related to the backpacks or otherwise. In our experience, most of the signs that could suggest ill-fitted tags occurred immediately after deploying and were addressed promptly. We found that most issues developed within the first two days of observation. Importantly, some issues were only detectable when birds were settled in their permanent housing environment where they could fly much more extensively. We also monitored the birds by assessing the tracking data to identify individuals that were outliers in the number of detections (suggesting they behaved differently to others). The main issue that arose after release into large aviaries was the backpack rubbing on the body or wings of the bird. Symptoms of this included bald spots on wings or neck, reduced movement or difficulty flying. These were addressed immediately by ensuring the backpack mount (and tray) were correctly fitted (i.e. not crooked and positioned away from the wings). However, in some cases, when the problem persisted, we completely removed the backpack, let the bird rest, and observed its behaviour without the backpack.

### Camera systems

Barcodes can be detected using either photos or video. The choice largely depends on the research question to be addressed, as well as the scale of data collection and its associated processing and storage requirements. In this section, we provide details on the necessary considerations for implementing a camera system, and details of our experience using several implementations, including high-resolution photos and video from action cameras, computer-controlled DSLR cameras, and the programmable camera module for the Raspberry Pi. We also discuss the pros and cons of each system for different types of research questions.

#### Code size and capture

For adequate detection and recognition of individual birds in photos or videos, the size of the barcodes should be at least 20 pixels per side in the captured image data (Crall *et al.* 2015), but this can vary depending on tag design and camera hardware. Detectability of tags can be improved by using high-resolution cameras, reducing the distances between the codes and the camera (either physically or by using zoom lenses), or increasing the physical size of the deployed tags (which is limited by the study organism). Other considerations such as lens distortion, sharpness, and depth of field must be considered depending on the setup and area being captured. Lens distortion can be partially corrected via software, but this correction reduces the effective resolution of the images, especially for wide-angle lenses (Fig. 3). Depth of field is an especially important consideration in situations where birds can perch at different heights. Finally, the camera shutter speed needs to be chosen carefully. Slow shutter speeds result in blurred, overexposed codes and thus failed detections. To prevent these problems, exposure time should be set as short as possible while ensuring that contrast and noise levels are adequate for the software to successfully read the codes. We found that darker images had greater detectability as they increased the clarity of the edges within the barcodes by reducing bleeding of the white areas of the barcode into the black areas.

#### Photos or video?

The choice between using photos or videos when capturing image data represents a classic tradeoff between spatial and temporal resolution. In general, video data offers higher temporal resolution by reducing spatial resolution, while photos offer higher spatial resolution by reducing temporal resolution. Both factors are limited by data throughput of the video hardware as well as data storage. Current imaging technologies vary widely in frame rates and image resolutions, and different camera setups can be adapted for data collection depending on the research question and project budget. Here we implemented and tested three types of recording devices:

*Action cameras*— We used GoPro Hero 4 action cameras to record video of the birds in a feeding arena 90×50 cm on the floor of the aviaries. We set the cameras to run continuously until the battery was depleted (approximately 45 minutes) and chose a resolution of 1920×1080 pixels (1080p) at 24 Hz to limit file size, reduce processing time, maximize battery life, and prevent the camera from overheating. We created a 3D-printed arm to attach the camera to the side of the cage, 50cm above the feeding arena and manually started recordings immediately after providing birds with a high-value food patch (to attract them to this focal area of the camera). A sample frame from a video of the feeding arena with codes detected is shown in Figure 3.

**Figure 3.**
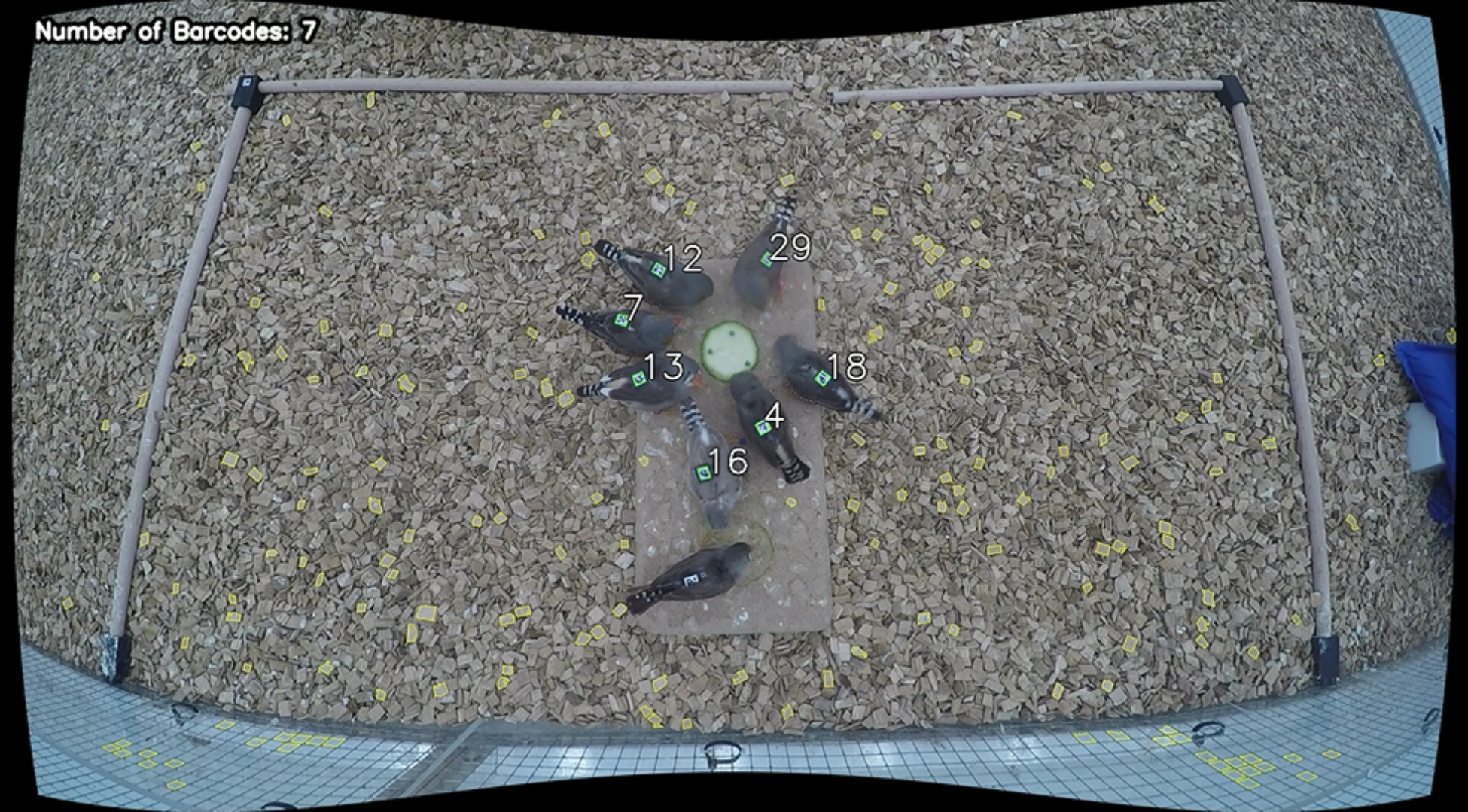
Barcode detections in a feeding context in one video frame (from a GoPro camera). The food source in the centre of the arena was pinned to a wooden board to prevent birds from moving it out of the frame. The pinpoint tracking algorithm (Graving 2017) detected barcodes on the back of each individual despite being on a complex, naturalistic background (wood chips). The yellow polygons are objects that were detected as candidate barcodes but did not match any known identities. The bird near the bottom of the image was not detected because its wings covered the barcode in this frame. Bird ID 16 was oriented away from the food. The black edges around image illustrate the software correction we used to partially compensate for wide-angle lens distortion.

These cameras produced adequate image quality but had noticeable distortion due to the wide-angle fixed lenses. We manipulated the resulting images to reduce distortion before running the detection code (see ‘extracting data from images’). At 1080p, we observed that the codes were sharp enough for detection, although the slow shutter speed at 24fps resulted in motion blur when birds moved. At 1080p resolution, we generated a 4GB file every 15-17 minutes of video, which is the maximum size supported by the cameras. This means that in a 45-minute recording session, we had to process three videos and store at least 12GB. Limitations of this setup include the need to manually operate the cameras, restricted recording time due to battery life or large file size, and a lack of options for automating the entire system (files had to be manually removed from the memory card and stored elsewhere). These cameras are capable of 4K quality video (2160p resolution), but this limits the recording time to 7-17 minutes, due to overheating.

*Digital SLR Cameras*—We briefly tested data collection using four Canon EOS1200 DSLR Cameras with 18-55mm lenses for recording video or still images.. We connected these cameras to Raspberry Pi 3 single-board computers to control the image capture frequency. We placed the cameras at the top of the aviaries facing directly down. The cameras were set to capture 10 frames at 1/200s every 10 minutes to measure the position of birds sitting on perches made from natural branches. These cameras can deliver high quality images up to 5184×3456 pixels (18 megapixels) and the zoom lenses allow for easy accommodation to different distances and to cover either small or large areas. However, because of the loud shutter, which visibly disturbed the birds in the enclosed aviary space, we abandoned this method. In video mode, it is possible to record high-resolution video (1080p) which is sufficient for collecting detailed movement data. Unfortunately, this mode can only record up to 30 minutes of video and must be started manually.

*Single-board Computers/Camera Modules*— We used Raspberry Pi 3 Model B with Camera Module V2 (RS Components Ltd and Allied Electronics Inc.) to record photos of birds on perches (Fig. 4). We installed two of these on top of each aviary, covering most of the perch system without overlap. We set the system to take 10 photos every 10 minutes to record the birds present at 1/200s shutter speeds. In our experience, one of the most important advantages of this system is the possibility of programming automation scripts via the picamera software package (Jones, 2013) in Python (Python Software Foundation, available at http://www.python.org). This approach gives the user fine-scale control over the quantity, sampling frequency, and spatial resolution of photos and videos. In combination with standard networking protocols like Secure Shell (SSH), these features allow for a fully-automated pipeline that includes image capture, file transfer, as well as processing and data storage when networked to a more powerful host computer. Another important advantage of these computers is their low cost, especially if the system requires multiple cameras per aviary or across multiple replicas in an experimental setup. Among the disadvantages of this system is the inconsistent quality of the camera modules (a small proportion of our cameras were unable to produce sharp images). To remedy this, we purchased additional cameras and chose the ones that produced the highest-quality images. Although these camera modules provide a large depth of field, they require manual focusing, which can be difficult and is often inconvenient.

**Figure 4.**
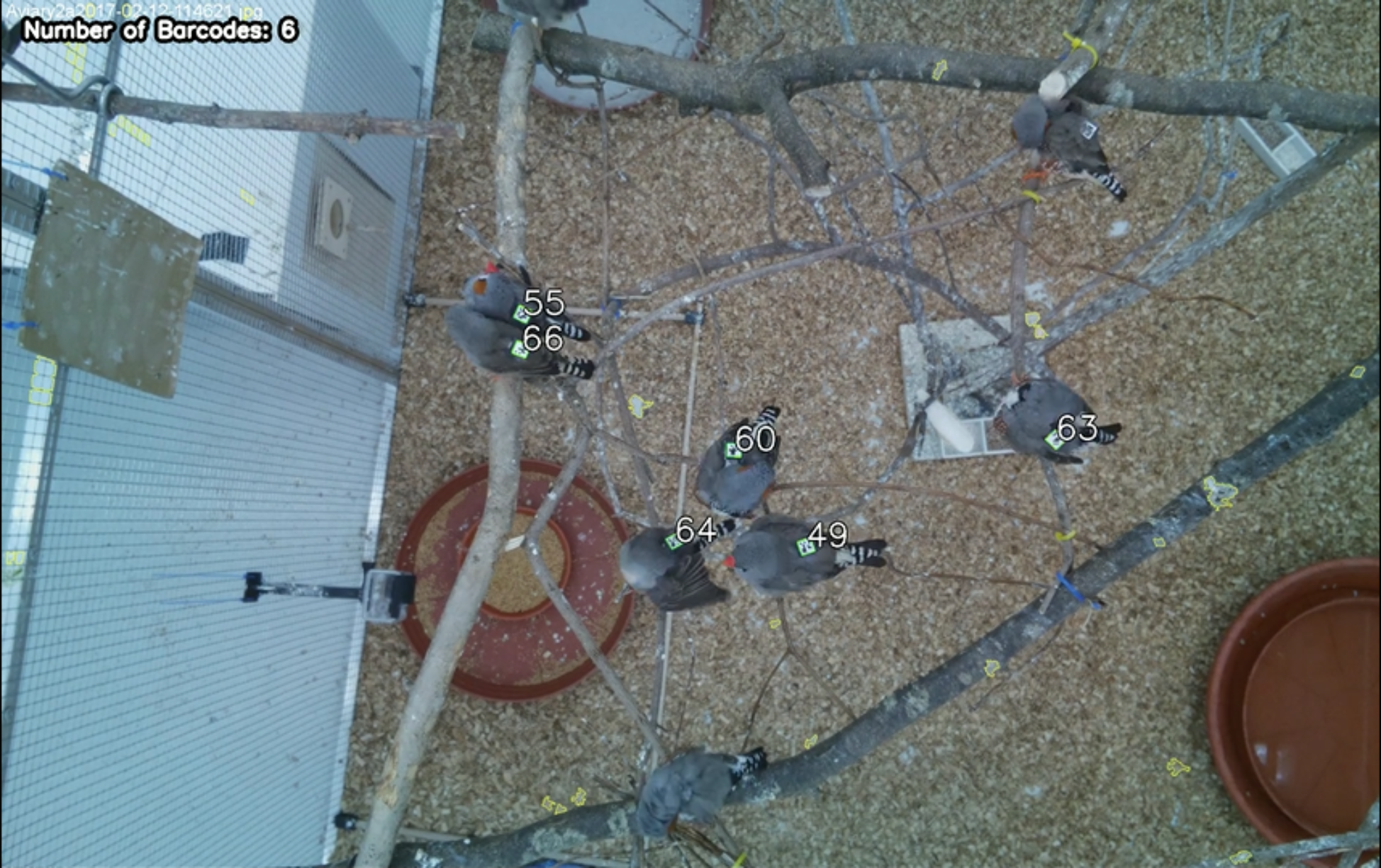
Barcode detections in a social perching context (photographed with the camera module of the Raspberry Pi). The software can easily detect visible codes in complex aviary environments and extract information about important interactions, such as direct body contact (individuals 55 and 66), that many tracking algorithms would fail to detect.

**Table 1:**
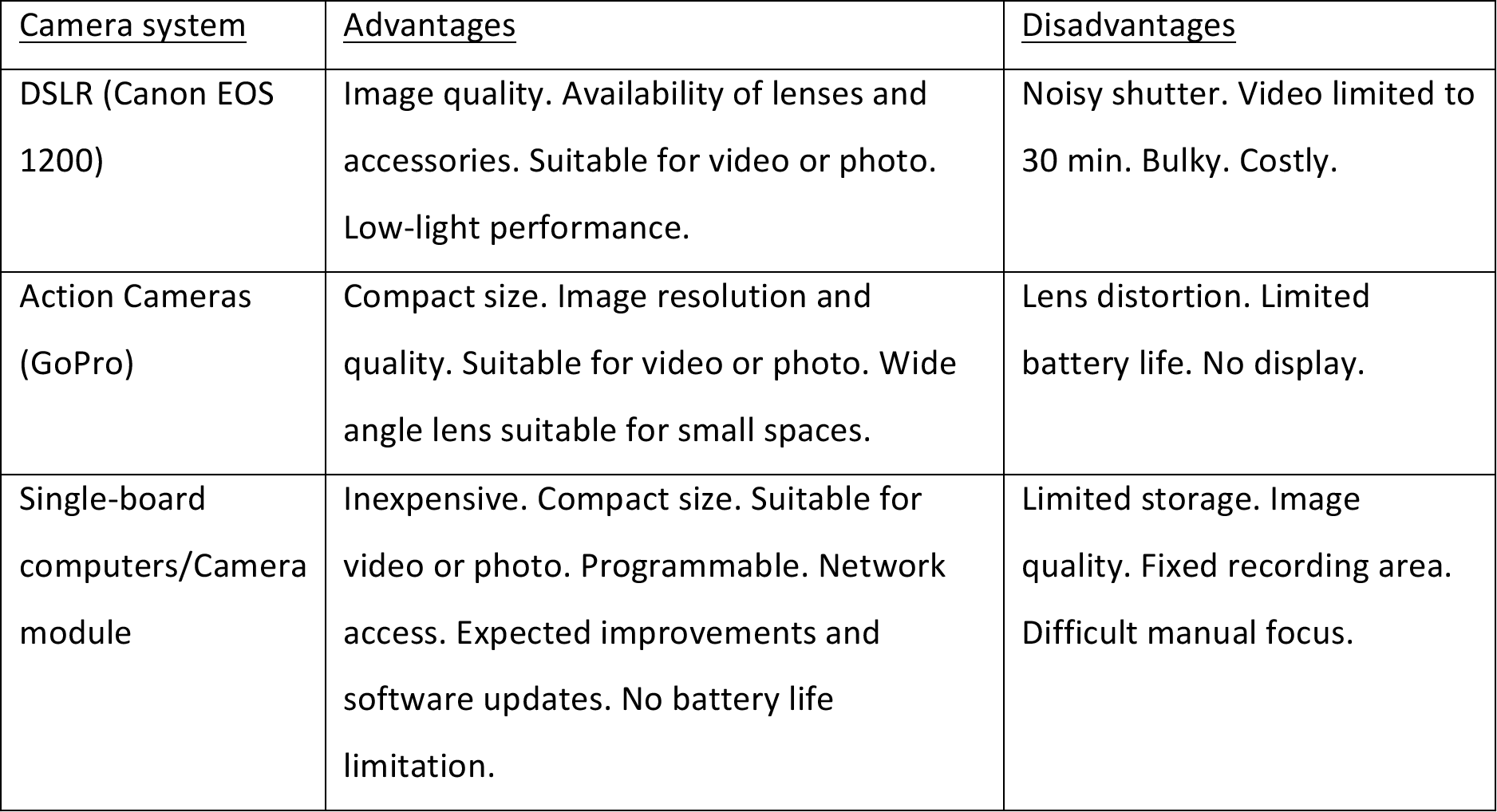
Summary of the pros and cons of different camera implementations tested.

### Extracting data from videos and images

Once videos or images are recorded, the next step is to extract location data from the barcodes contained in the image data. Several libraries are available to accomplish this (Crall *et al.* 2015; Graving 2017), and each provides its own set of barcodes. The libraries extract the tag identity and locations of the internal corners for each detected code, which can then be used to calculate the position and orientation of individuals. In our study, we used the software library pinpoint by Graving (2017).

#### Code Detection

The detection algorithm finds the identity matrix of the barcode using the contrasting white and black edges between the barcode and the black frame of the plastic tray on the backpack. Images are binarised using an adaptive (spatially-localised) thresholding algorithm (which allows for uneven lighting) and candidate barcodes are detected based on their geometry (which allows for complex backgrounds). Once a candidate barcode is detected, the identity matrix is extracted from the pixel data and compared to known identities stored in a tag dictionary. The pinpoint tracking algorithm (Graving 2017) can reliably detect the codes at any arbitrary angle and even when they are not completely perpendicular to the central-axis of the camera lens. The software provides the identity of each tag and Cartesian coordinates (relative to the top-left of the frame) for the corners of each detected barcode with sub-pixel resolution, which can be used to calculate the orientation of the code (note the importance of fitting the code in the right direction on the birds).

#### Interpolation of individual positions

Since quick movements and changes in body position may affect the detection of individuals as they hop around the feeding arena, we found that we could use linear interpolation to fill gaps between detected positions if those detections were less than 1 second apart (24 video frames of video when using the action camera). We calculated the length and orientation of the movement between the two detections. The average distance moved for each missing frame was calculated by dividing the distance moved between detections by the total number of frames being interpolated, and for each missing frame, the individual’s position was shifted along the vector between the two points by the average distance. This interpolation method was not possible when using photos.

### Example data analyses

To demonstrate the use of this automated approach to data collection and analysis, we studied the foraging behaviour of individual zebra finches at a high-quality food source and constructed foraging networks based on high-resolution movement data measured using our system. Social networks are particularly challenging to study using manual observation because they require measuring the behaviour of most or all individuals simultaneously. To achieve this, we created an arena 90 × 50 cm on the floor of each of the two aviaries and provided birds with an ephemeral high-quality food resource (a slice of zucchini/courgette) twice per day (around 9am and 4pm). We used a barcode to record the centroid of the resource, which was subsequently removed to allow birds unobstructed access to the food. Birds were fasted for an hour before the experiment to ensure they were motivated to feed, and their access to the food resource was captured on video using the GoPro Hero 4 camera fitted 50 cm above the food (see above). We collected data on the two aviaries during 58 days, between December 15 2016 and March 29.

We extracted feeding association data, representing the propensity for individuals to synchronise their feeding and tolerate one another at the food source. We recorded the identity of each individual detected at the food for every video frame by defining a feeding zone with respect to the centroid of the food resource. A feeding event was recorded when a barcode was detected within a 154-pixel (or approximately 8-cm) radius of the resource centroid, and the bird was facing the food (i.e. the centroid was within the 180° zone in front of the bird)(Fig. 3). Once we identified the individuals in every frame and classified feeding events, we constructed a weighted, undirected social network representing the co-feeding relationships among individuals (represented as nodes) in each flock. We accomplished this by transforming our data into a matrix of pairwise associations using a simple ratio index (*SRI*) for every pair of individuals in each flock. Here, the edge weight between two individuals (*SRI*_*ij*_) is the probability of observing individuals *i* and *j* feeding together given that either *i* or *j* has been detected. When using images, this calculation is simply given by:

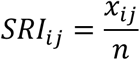

for *i*, *j* = 1,…*N* and *i* ≠ *j*, where *N* is the total number of individuals in the flock, *x*_*ij*_is the number of frames in which individuals *i* and *j* were feeding together, and *n* is the total number of frames where either *i* or *j* was detected (alone or together).

## RESULTS

### Barcode deployment and maintenance

We deployed backpacks on 58 zebra finches (Fig. 5), which required about three minutes of handling for each individual, plus observation and monitoring time. All the deployed backpacks lasted throughout the experimental period without causing any injuries to the birds. However, minor maintenance was required as backpacks and codes showed some wearing due to grooming and allopreening (see backpack-mount in Figure 5). Common issues included loss of ink on and around the barcodes, weakened paper around the front holes, and unglued mounts. We also noticed that, in a few cases, the straps lost elasticity after four months and appeared loose. More commonly, we observed that debris (i.e. food remains or excrement) on the barcode obstructed its detection. Every time we detected one of these issues, we addressed it immediately to guarantee both safety of the birds and quality and continuity of the data collection. For any minor issues, we carefully cleaned the codes to remove debris, or covered the ink-less spots with black ink permanent markers. For extensive damage on the mount or the surface of the barcode, we removed and replaced the mount keeping the strip and the elastic string on the bird, thus reducing manipulation and acclimation time. In cases that required a whole new backpack, we repeated the process of the first deployment.

**Figure 5.**
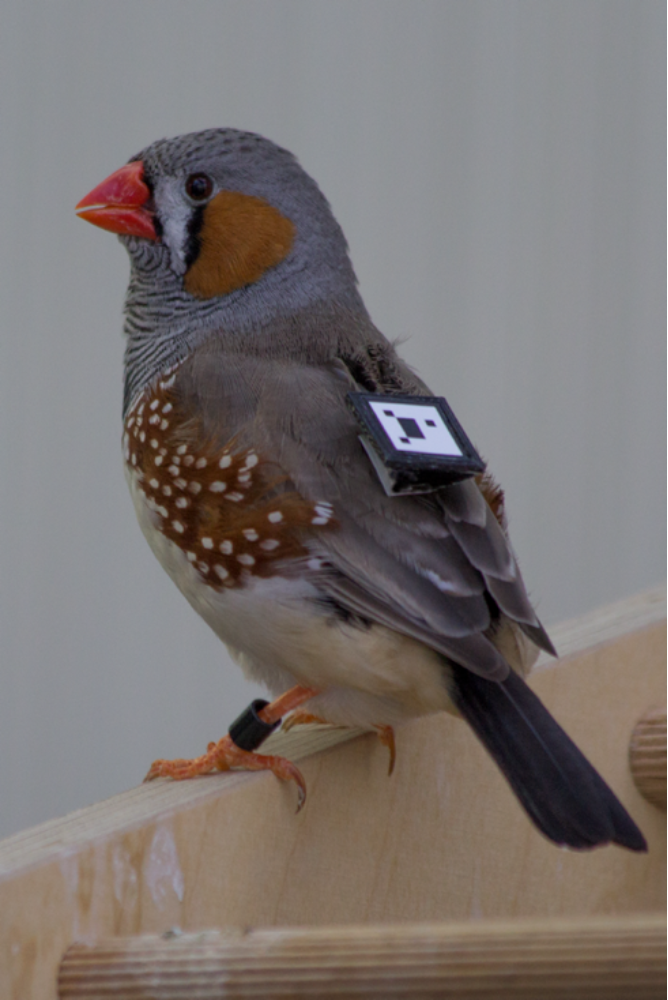
Male zebra finch with backpack and barcode approximately 4 months after fitting. Note the wearing on the backpack structure caused by grooming behaviours.

### Detection

We recorded 48 hours of video at the feeding arena using GoPro Cameras and recorded photos over 6 days using Raspberry Pi cameras. The detection software identified 52.05% of the barcodes (i.e. birds) present in 100 randomly-selected frames from the GoPro footage. This percentage was improved to 64.58% after linear interpolation. The software detected 60.40% of the birds present in 100 randomly-sampled images captured using the Raspberry Pi cameras. The most common reasons for non-detections were motion blur and feathers temporarily obscuring parts of the code (e.g. Fig. 3). However, we note that, even at 1 second resolution, barcodes were detected on average 6 out of every 10 seconds, which should be sufficient for the vast majority of applications. Detection rates can be considerably improved by trimming feathers around the code and optimising the camera setups.

### Example data analyses

Using image data collected with a GoPro mounted over the food arena, we were able to distinguish birds consuming the resource from those present in the frame but not feeding (Figure 6). Further, from the single 45 minute observation period shown in Figure 6, we recorded 74960 records of individual positions. These records also contain many potential interactions. We demonstrate that the data on the co-presence of individuals at a food source can be used to generate social networks (Figure 7), a powerful approach used in many studies of animal behaviour for which extensive observation data are required.

**Figure 6.**
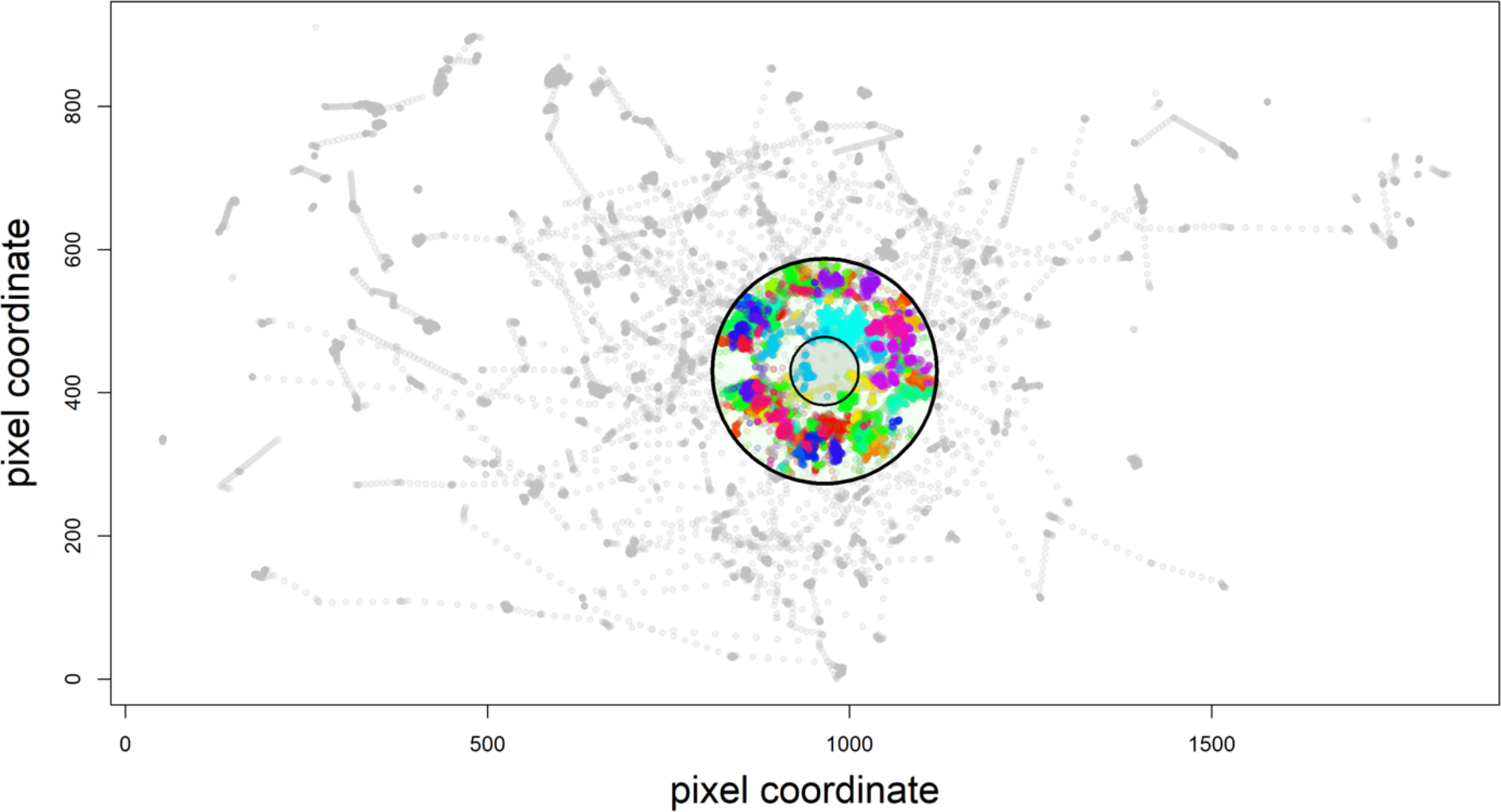
Detection of feeding behaviour characterised by proximity and directionality to the food source. The inner circle represents the outline of the food source, and the outer circle represents the 154 pixel boundary for birds to considered to be ‘at food’. Coloured dots represent detections of different individuals within the ‘at food’ zone. Grey dots are birds present and identified in the frame but not actively feeding. The positions of birds away from the centre of the frame are less accurate due to lens distortion (See Figure 3).

**Figure 7.**
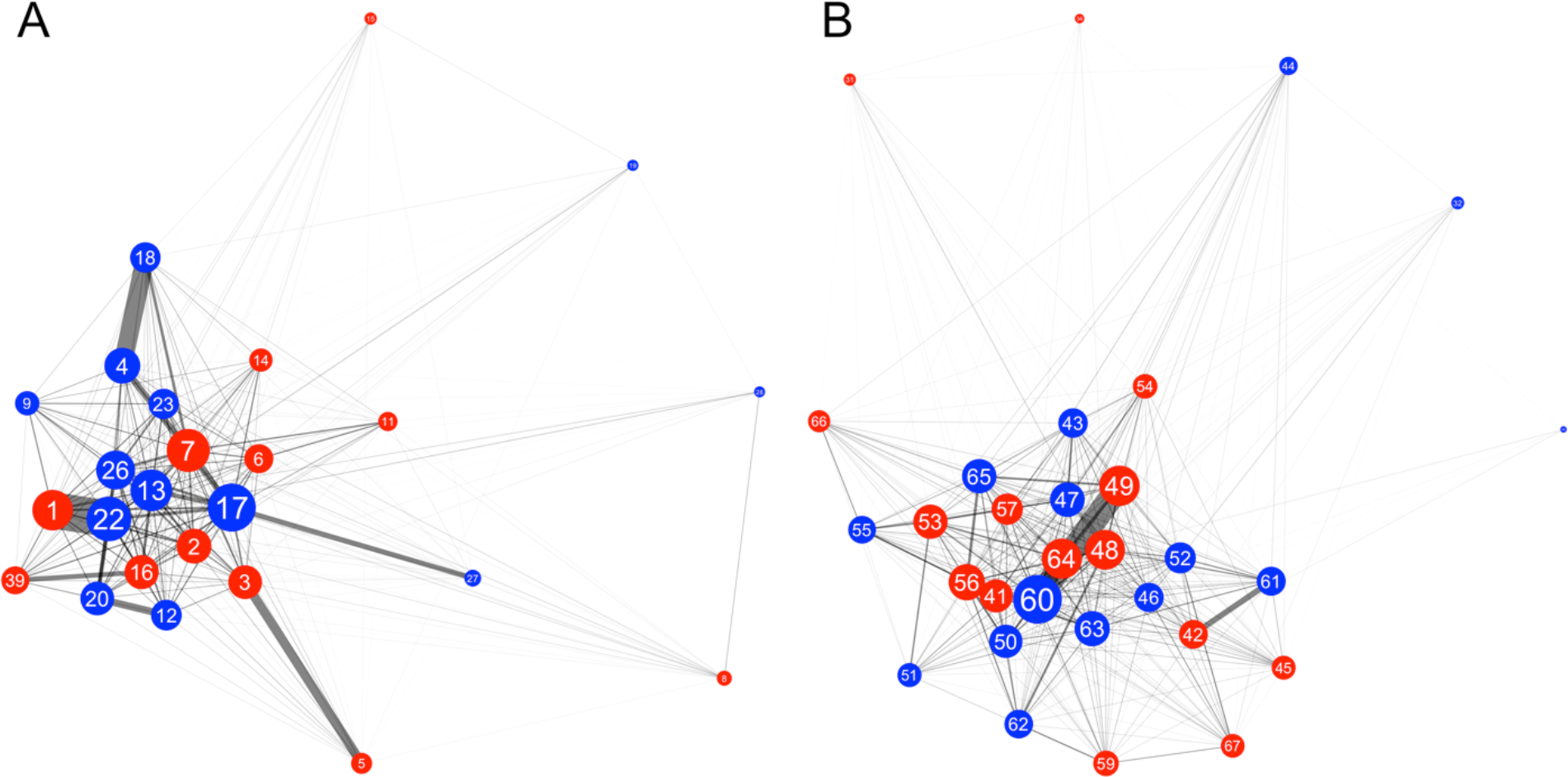
Affiliative networks generated with co-feeding data extracted from barcodes detected at a food source using a video camera in (A) flock 1 and (B) flock 2. Each node (circle) represents an individual. Red nodes represent males, and blue nodes represent females. The size of the node represents the individual’s degree in the network, a measure of centrality computed by summing the weights of all the edges connected to it. The thickness of the line represents the strength of the association between each pair of individuals.

## DISCUSSION

We present a method to automatically measure the behaviour of captive birds using backpack-mounted barcodes, image capture, and computer detection. With proper deployment, manipulation and monitoring, we have shown that this system is safe for the birds, durable and capable of delivering extensive data on individual identification of subjects, including their position and direction. This system presents several advantages to more commonly-implemented methods. In particular, it is adaptable to different contexts and research questions, being possible to vary the temporal resolution (photos or video) and the area covered, without requiring any additional markers to birds. For general purposes, the use of Raspberry Pi single-board computers and camera modules makes this method affordable, enabling high-throughput data collection over multiple samples and subsequently increasing sample size and statistical power. Our example analyses demonstrate that the barcode-based approach can generate similar data to what is often collected using PIT tags (Fig. 7), but also provides much richer information on movements and spatial location within patches (Fig. 6). We found that the backpack system simplified the data analysis because we were certain about the co-occurrence of birds at the same food source (i.e. captured in the same frame), instead of having to infer co-occurrences from sequences of detections using pattern-recognition algorithms (e.g. Psorakis *et al.* 2015).

We tested the application of different camera setups and behavioural contexts, including video for feeding arenas and photos in co-perching scenarios. We found this system is easily adaptable and that cameras could be fitted above bathing areas, outside of nest boxes, and potentially in open areas to capture birds in flight. The latter is feasible with shutter speeds above 1/1000s, which is possible with the Raspberry Pi camera modules. The decision the type of camera and on video or photos will depend on each research question. For example, researchers could choose video for recording dominance interactions or other interactions that involve movement, or capture photos every few seconds to capture affiliative data, which are useful in the study of social networks, pair formation, or group stability. The type of data provided by these barcodes, combined with detection software like pinpoint (Graving 2017), provides new opportunities for analysis. Using machine learning, it will be possible to automatically classify behaviours and interactions over extended periods of time while also minimizing manual annotation by a human observer (Robie *et al.* 2017), thereby avoiding bias and fatigue. Such approaches have been developed for studying other organisms (Kabra *et al.* 2013), which use data that is similar to what our system generates.

Our backpack-based barcode method has potential to be adapted to diverse range of systems or to be enhanced with additional equipment, full remote access, or other accessories as required to address behavioural questions. Although we only collected data during daylight hours, barcodes could easily be detected in low-light conditions and many commercially-available infrared cameras can image the black-and-white codes without visible light. While most birds are not very active at night, there is increasing evidence that many important behaviours happen early in the morning (Bonter *et al.* 2013). Such behaviours could easily be captured with this barcode system but would be almost impossible to study using manual observations or video as it is difficult to identify coloured leg bands. Future applications include using barcodes to identify individuals interacting with a device (e.g. a feeder or a puzzle box). To date this has mostly relied on using PIT tags (e.g. Aplin *et al.* 2015), which limits sampling to a single individual at once. In social species, individuals often congregate, and a barcode system can facilitate multiple simultaneous detections and quantify relative positions of individuals to one-another and to the device. The implementation of ‘real-time’ detection could allow for algorithms that control devices in response to the behaviour of birds, such as allowing only a maximum number of individuals in one area or selectively dispensing food to particular individuals (as performed by Firth *et al.* 2015). Barcodes could provide a powerful interface between individuals and experimental devices, not only by being able to provide tailored responses (such as individual learning algorithms, Morand-Ferron *et al.* 2015), but also, unlike almost any other system, by capturing information about who else is present when particular events occur.

Although we have discussed the multiple advantages, the limitations of the system must be also considered. While backpacks and barcodes can last for more than four months, permanent monitoring was required to assure safety of the birds and adequate delivery of data. Grooming and allopreening caused some wear on the backpacks and codes, and this sometimes led to impaired movement of the birds. Detecting and addressing such issues is important for both safety of the birds and continuity of the data collection. Additionally, there are unavoidable issues that reduce detectability, like fast movement, codes tilted due to extreme body position, and wings or feathers partially covering the trays. The current design of the backpacks addresses these issues well and delivers consistent detection (which can be improved using linear interpolation when using video). Additional concerns related to camera systems, such as storage, resolution, lens distortion or lighting, can be solved for specific research circumstances.

A key question that requires further investigation is whether these backpacks will be suitable for field deployment. We found that in zebra finches, we could detect most issues within the first 1-2 days. However, few field studies are amenable to keeping birds in captivity to allow such monitoring. Thus, field applications may be limited to species that either have well-established protocols for fitting backpacks in the field or those in which individuals can be easily monitored (e.g. territorial species). We believe that there is a danger that small songbirds could entangle their backpacks in small branches, particularly if backpacks become loose over time. Finally, our aviaries had artificial lighting that remained constant during daytime. Natural lighting conditions for outdoor studies must consider the changing environment (i.e. sun position and cloud coverage) to avoid unusable images due to the differences in light quality from dawn/dusk to noon. For example, sun shining directly on the white tag will make the code invisible to the camera, while a setup designed for sunny conditions would create completely black photos under cloudy conditions. The use of infra-red cameras and infra-red lighting is one way to overcome this challenge.

Our backpack-mounted barcode system could revolutionise data collection in a range of experimental systems. We have demonstrated that it can be implemented safely and cheaply. Further, it has the ability to collect extensive data across many individuals simultaneously and the flexibility to address diverse research questions. With simple software modifications, the system can also be integrated into active devices that interface directly with individuals, which will prove to be an extremely powerful experimental approach.

## Supplemental Data

A sheet containing the backpack template is provided in Supplemental Materials 1.

## Acknowledgments

We thank Jana Hörsch, Laila Darouich, Alex Bruttel, Daniel Zuñiga, and the animal caretaker team at the Max Planck Institute for Ornithology in Radolfzell for assistance with data collection and monitoring animal health each day. We also thank Iain Couzin for his support of the project. This work was funded by the Max Planck Society.

